# The effects of estrogen exposure on survival, growth, and fecundity of *Daphnia magna*

**DOI:** 10.64898/2026.06.27.734946

**Authors:** Siobhan J Boyle, S Schaack

## Abstract

High concentrations of steroidal hormone compounds are a growing source of concern for environmental pollution in aquatic ecosystems. In this study, we examine the effects of two estrogenic compounds (estriol and 17α-ethinylestradiol) on fitness traits in the aquatic microcrustacean, *Daphnia magna*, a key bioindicator species for toxicology studies. The impacts were compared of two forms representing a natural and synthetic estrogenic compound. Growth and reproduction traits were assayed by exposing *Daphnia* to each estrogen type at four concentrations reflecting potential environmental exposure conditions up to acute toxicity levels (ranging from 0.1 - 50 µg/L). Assaying the effects at a variety of concentrations is important given that it is known that hormone exposures can often result in non-monotonic responses. Both forms of estrogen impact a subset of the traits assessed, in some cases leading to beneficial changes and others causing harm. Estriol, the naturally-occurring estrogen, and EE2, the synthetic version, at high doses shift fitness traits in opposite directions such as adult growth rate as do at low doses for fecundity. In conclusion, our results support the need to assay a wide array of traits using multiple forms of steroidal hormones at a range of doses in order to assess non-monotonic patterns and their impact on an organismal fitness. In particular, assays that extend beyond the conventional measurements of lethality during acute exposure windows will be essential for understanding the impact of increased levels of hormone pollution on aquatic organisms and ecosystem health.

## Introduction

### Hormones in the Environment

Hormone and pharmaceutical contamination is a topic of growing concern with the increased number of studies showing effects of endocrine-disrupting compounds (EDCs; e.g., Nobre et al. 2024). Both naturally-occurring and synthetic hormones are known to impact ecosystems when released into the environment causing feminization, reproductive impacts, and even DNA damage (e.g., Paravani et al. 2024; Sauvé and Desrosiers 2014; Yan et al. 2012; Viglino et al. 2008; reviewed in Ojoghoro et al. 2021). Estrogenic hormones, specifically, may pose a threat to human and environmental health (De Aquino et al. 2021), although assays of specific fitness effects resulting from exposure are more limited (e.g., Luna et al. 2015 and Rodrigues et al. 2021).

While naturally-occurring estrogens can be released into the environment by organisms, synthetic counterparts used in birth control and hormone therapies are often introduced into wastewater via medical, agricultural, and industrial discharge, in some cases at high concentration (Tang et al. 2021; Fick et al. 2009). The presence of estrogenic contaminants has been reported worldwide in surface, ground, ocean, and even drinking waters, often at low concentrations (less than 0.1 ng/L) but as high as >10,000,000 ng/L (Brossa et al. 2005; Aris et al. 2014; Rodrigues et al. 2021; Da Silva et al. 2025; Almeida et al. 2020). Even at low concentrations, hormonal pollutants can adversely affect reproductive and developmental processes in humans and wildlife (Zhang et al. 2016; Jackson et al. 2020; Dietrich et al. 2010). Predicted no-effect concentrations (PNECs) have been estimated for many estrogenic compounds, however they are frequently exceeded in environmental samples (Unnikrishan et al. 2024; Caldwell et al. 2012). Current data suggests that naturally occurring estrogens have higher PNECs than the synthetic counterparts, and are seemingly less impactful in small dosages (Caldwell et al. 2012; Huber et al. 2004). EDCs and hormones can have non-monotonic, or even unpredictable response curves, where exposure to high doses does not predict low dose effects (Vandenberg et al. 2012).

### Structure and Properties of 17ɑ-ethinylestradiol and Estriol

The many forms of estrogenic hormones, natural and synthetic, can vary in their metabolic function and their ability to degrade (Lai et al. 2008, Müller et al. 2021). In general, steroidal estrogens share a tetracyclic molecular structure, with a stable aromatic A ring and phenolic hydroxyl group (Adeel et al. 2017; Okkerman et al. 2001; Figure 1). They are hydrophobic with high molecular weights, resulting in longer degradation time and low solubility although the half-life depends on oxygen availability, environmental conditions, and the presence of certain bacteria (Pauwels et al. 2008). Estrogens are often hydrolyzed or methylated by the liver and conjugated with glucuronic acid or sulfate before being excreted (Souza et al. 2013; Denver et al. 2019; Stanczyk 2024). Naturally-occurring estrogenic hormones originate primarily from human and animal byproducts in partially metabolized or unmetabolized forms (Bilal et al. 2021; Zhao et al. 2019), the most common of which are estrone, estriol, and estradiol. Estriol is commonly used in menopausal hormone therapies and is also produced and excreted in large quantities by pregnant women (Ali et al. 2017).

**Figure 1.**
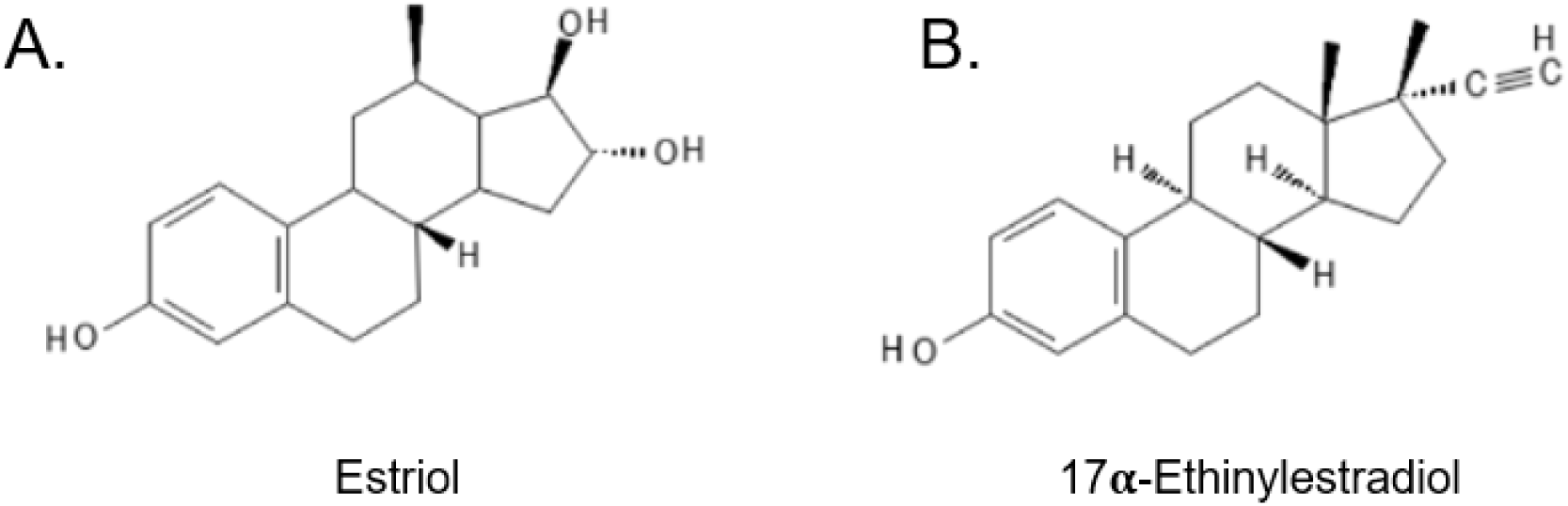
Chemical structures of two forms of estrogenic hormones used in this study: (A) estriol, a naturally-occurring form of estrogen and (B) 17α-ethinylestradiol (also referred to as ‘EE2’), a common synthetic form of estrogen found in birth control and other hormone therapies.

A common synthetic estrogen, 17α-ethinylestradiol (referred to as EE2), is frequently used in contraceptives due to its ability to interact with receptors and mimic natural estrogens (Baatrup and Henriksen 2015; Rodgers-Gray et al. 2000). EE2 undergoes less metabolic transformation and is excreted as an unchanged molecule 80% of the time (Thomas and Potter 2013). EE2 is an alkyne compound containing an ethinyl group on the 17th carbon responsible for stabilizing the compound and increasing resistance to oxidation (Payus et al. 2021; Feng et al. 2010). EE2 is known to be light sensitive and can undergo photolysis, but studies have shown it is more persistent in groundwater systems compared to naturally-occurring estrogens (Pal et al. 2010; Tapiero et al. 2002). EE2 has been heavily investigated since it was identified as a pollutant of emerging concern with hormonal and steroidal effects (Tiedeken et al. 2017; Valbonesi et al. 2021; Li et al. 2023; Hultman et al. 2015; Ying et al. 2002). Compared to EE2, estriol lacks the triple bonded carbon group and instead contains two tertiary alcohol (OH) groups (Figure 1). The structure of both estrogens allows for them to interact with a wide variety of hormone and steroid receptors, generating high endocrine-disrupting potential (Pamplona-Silva et al. 2018; Kernen et al. 2022; Gutendorf and Westendorf, 2001). Studies directly comparing exposure effects of a naturally-occurring (estriol) and synthetic (EE2) version of estrogen (e.g., Liarte et al. 2011) are still needed.

### Effects of estrogens on aquatic organisms

Observations of hormone contamination on wildlife have been published for decades (e.g., Burns et al. 1984), but controlled experimental assays are crucial for understanding of the effects (Kashian and Dodson 2004; Da Silva et al. 2025). Early studies suggested potential structural similarities between human estrogens and invertebrate prohormones and emphasized a high level of conservation between endocrine-related processes across vertebrates and invertebrates (Leblanc and Mclachlan 1999; Warrier et al. 2001; Pertseva and Shpakov 2002). Estrogen contamination has been studied in many freshwater organisms, including fish, mollusks, insects, and shrimp (i.e., Salla et al. 2024; Mitsui et al. 2007; Bovier et al. 2018; Islam et al. 2020; Hamilton et al. 2022; Leroux et al. 2025). In many cases, thresholds or effects have differed based on dosage or the specific type of estrogenic compound used for the exposure. For example, in *Danio rerio*, EE2 was found to inhibit reproduction at concentrations as low as 9.3 ng/L and decrease social behavior, even just above the predicted no-effect concentration of 0.1 ng/L (PNEC; Tan et al. 2024; Porseryd et al. 2017; Schäfers et al. 2007; Conley et al. 2017; Caldwell et al. 2012). In contrast, studies using estriol found that a concentration of 21.7 µg/L was needed to influence sex ratios in zebrafish (Holbech et al. 2006). Similarly, a number of studies in other fish (*Fundulus heteroclitus* and *Oryzias latipes*) performed using a range of concentrations and with different estrogen types show dosage-dependent effects on a wide variety of traits (MacLatchy et al. 2003; Tilton et al. 2005; Hogan et al. 2010; Lei et al. 2014). Such studies argue for the importance of investigating the effects of estrogen exposure on organismal fitness and comparing estrogen types across concentrations in a single experiment (e.g., Vaillant et al. 2020; Liarte et al. 2011).

In this experiment, *Daphnia magna* were exposed to EE2 and estriol at four dosage levels (0.1 µg/L, 1 µg/L, 10 µg/L, and 50 µg/L) and compared them to control conditions. This range of concentrations reflects potential environmental conditions up to an acute exposure level (Djebbi et al. 2022; Svigruha et al. 2021). *Daphnia magna* is a commonly used model organism in ecotoxicology, often used to assay ‘no effect’ or ‘lethal count 50’ concentrations of pharmaceutical pollutants, which are the levels at which no harmful effects or 50% mortality are observed, respectively (Ebert 2005). *Daphnia* are, however, also very well studied in other biological subdisciplines and have a number of readily quantifiable, sublethal traits that can be linked to the health of individuals or of whole ecosystems, given the roles *Daphnia* play in food webs as primary consumers (Pinos-Vélez et al. 2023; Ebert 2022; Jaser et al. 2003). Environmental studies on invertebrates like *D. magna* are often overlooked as there is a lack of clear connection between conserved vertebrate hormones and invertebrates (Baldwin et al. 1995; Goto and Hiromi, 2003; Segner et al. 2003).

## Methods

### Experimental protocol

*Daphnia magna* used in this experiment were all derived from the same ancestral genotype (referred to as genotype ‘GC’ in previous studies; see Ho et al. 2021) and have been reared in lab conditions since being obtained from Dieter Ebert in 2014. For this experiment, animals were kept in an environmental chamber at a constant temperature of 19 ℃ under a 12:12 light:dark cycle. Media was renewed every 1-2 days for all animals in the experiment and animals were fed a constant amount of algae (*Scenedesmus obliquus*) *ad libitum*. Females were monitored daily and neonates were introduced into treatment and control conditions at <24 hours old (Hamilton et al. 2022). For the first 5 days of the treatment period, neonates were maintained in low density groups of 5 to 6 individuals in 40 mL of media (Aachener Daphnien Medium [ADaM]; Klüttgen et al. 1994) in 50 mL glass tubes. After five days, individuals were moved to new glass tubes and housed and monitored individually for the remainder of the experimental period.

Estrogen exposures were performed using stock solutions made at 1 mg/L concentration using estriol (Cayman Chemical Company, Item no. 10006484, ≥95%) and 17α-ethinylestradiol (EE2; Sigma-Aldrich, E4876, ≥98%) dissolved into deionized water. Stock solutions were made weekly, and were used to make each concentration level by adding the appropriate amount of the stock solution to ADaM. Control conditions were ADaM only. Individuals were checked daily for visible signs of maturity (visible eggs in brood chamber), any released clutches, and death. Length measurements were taken on day 7 and 21 using an Olympus-SZ2 microscope (40x) calibrating the length using the eye piece graticule. Lengths were log_10_ transformed prior to analysis to account for allometric scaling when body sizes differ.

After releasing their third clutch, typically after 28 days of exposure, oxygen consumption was assayed using a PreSens SensorDish™ plate reader. Individuals were acclimated to room temperature (∼20 ℃) overnight. Individuals were removed from the treatment media and allowed to swim in sterile (autoclaved) ADaM for 15 sec, and then transferred into a vial filled with sterile ADaM. Vials were sealed and placed on the SensorDish reader, which was kept in a dark constant temperature chamber. Oxygen consumption was measured every 3 min for 3 hours for each individual. Individual body length was measured and dry mass was recorded after drying animals at 60 ℃ for 2 days.

### Statistical Analyses

All analyses were performed using RStudio (version 4.5.1; Posit Team 2025). The code used to perform the analysis are available in Supplemental Files S1. All data are available in Supplemental Table S1. After data visualization and model testing (Supplemental Table S2), estrogen types were treated as separate factors and dosage levels as categorical variables given the non-monotonic and diverse patterns observed across the eleven measured traits. Survivorship was analyzed using Cox Proportional Hazard tests. For data related to growth and fecundity traits, in some cases, models with many parameters offered the best fit across dosages (see Supplemental Table S2). Plots of the data illustrate there is high variance among treatment and dosage levels across traits (Figure 3 and 4), making us comfortable with the violation of the independence assumption (among concentrations) inherent to using analysis of variance (ANOVA). ANOVAs were performed, treating concentration levels as independent and using post-hoc Tukey tests to further examine differences among dosage levels (as in Doyle et al. 2013). Post-hoc tests were used to test differences between a) dosage levels within each estrogen type, b) estrogen types at each dosage level, and c) treatments to controls (dosage = 0).

## Results and Discussion

*D. magna* were exposed to two types of estrogen (EE2 and estriol) at a range of concentrations (0.1, 1, 10, and 50 µg/L) in order to assess the impact of treatment, dose, and interaction effects on a variety of fitness traits. Hereon, the term ‘estrogen’ is used to refer to results from exposure to either estriol or EE2, although in most cases results are presented for each compound separately and type is specified. We assayed survivorship (Figure 2 and Table 1) in all animals introduced into the experiment, and in animals that survived to adulthood, we measured growth and oxygen consumption (Table 2, Figure 3, and Supplemental Table S3), in addition to fecundity traits (Table 3, Figure 4, and Supplemental Table S4).

**Figure 2.**
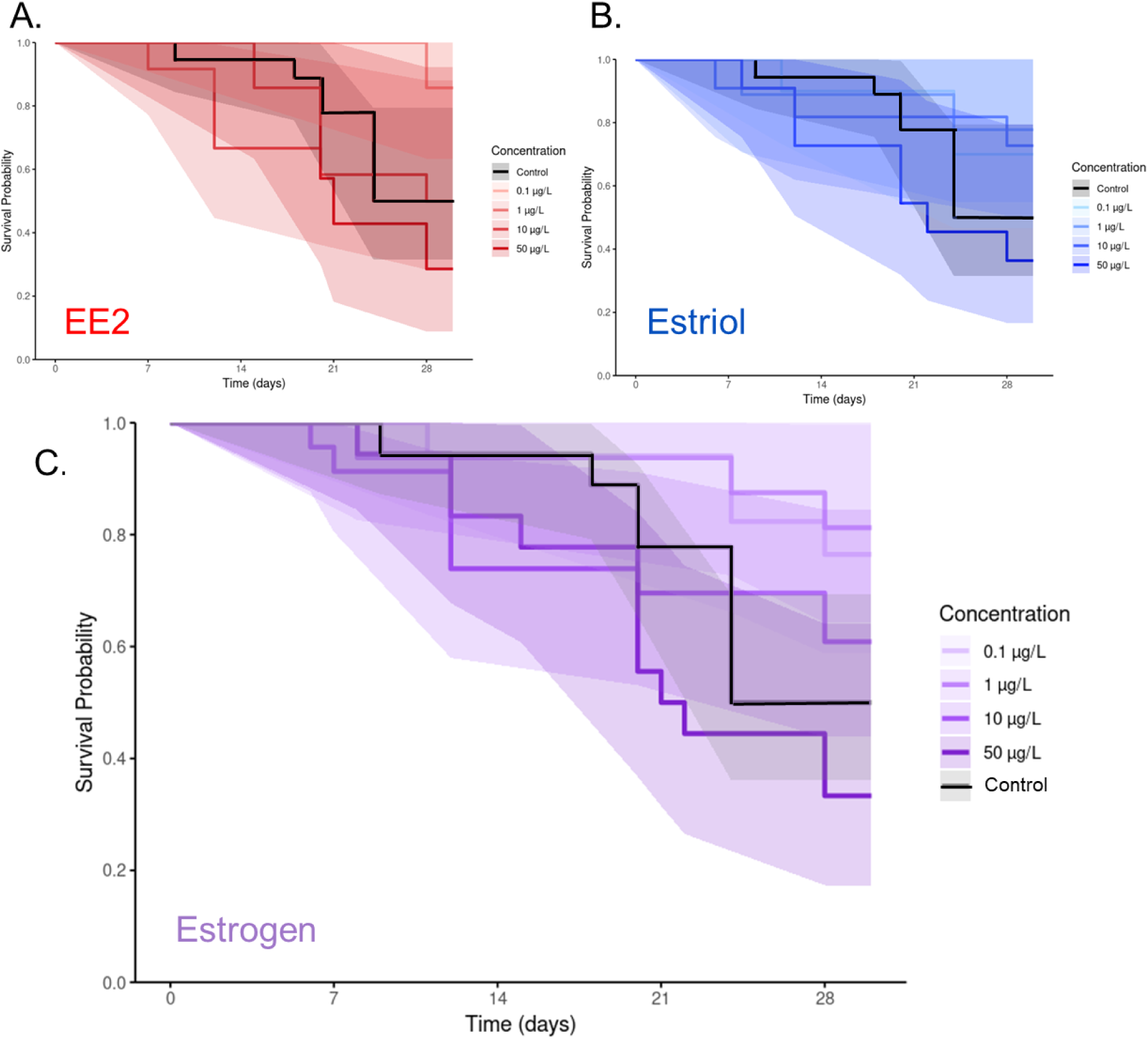
Survival curves depicting the probability of survival over time for each dosage with controls (no hormone) shown in black for (A) individuals exposed to 17α-ethinylestradiol (EE2) in red, (B) individuals exposed to estriol in blue, and (C) pooled data for animals exposed to either type of estrogen in purple. Higher concentrations are depicted with darker lines in each case, and shaded areas represent 95% confidence intervals around mean values at each time point.

**Table 1.**
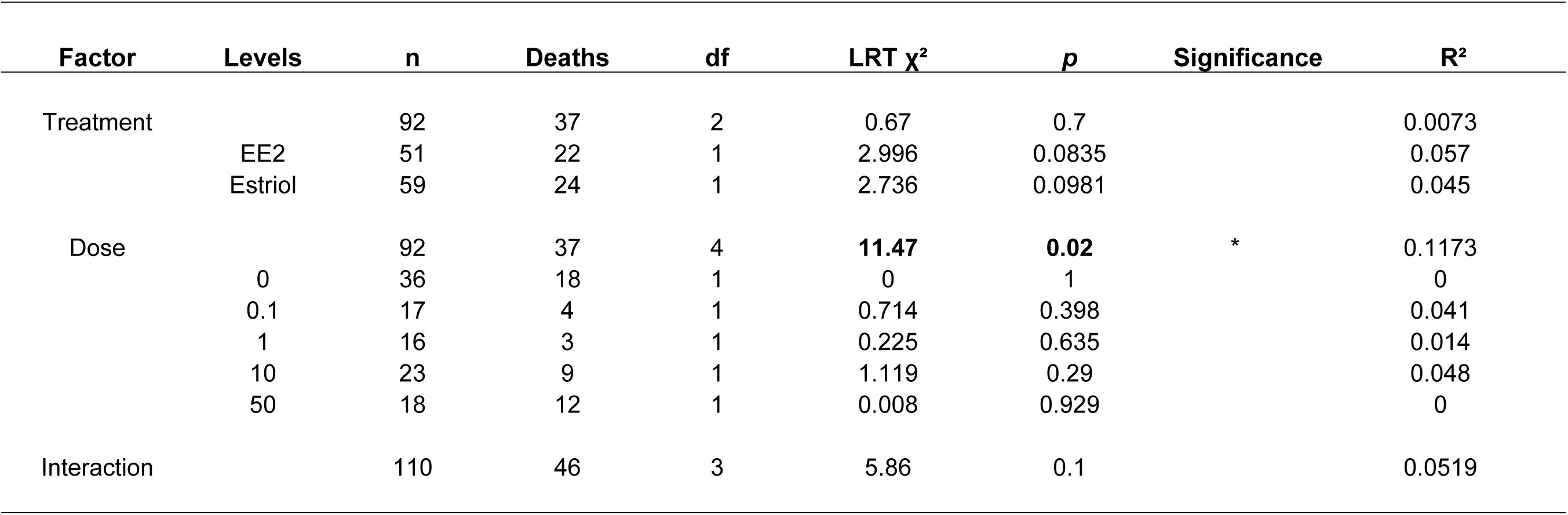
Effects of estrogenic compounds (estriol and EE2) across a range of doses (0, 0.1 µg/L, 1 µg/L, 10 µg/L, and 50 µg/L) on survivorship for *D. magna* (LRT = likelihood ratio test; * denotes *p* < 0.05, ** denotes *p* < 0.01, *** denotes *p* < 0.001).

**Table 2.** Effects of estrogenic compounds (estriol and EE2) across a range of doses (0, 0.1 µg/L, 1 µg/L, 10 µg/L, and 50 µg/L) on body size, growth, and oxygen concumption for *D. magna* (* denotes *p* < 0.05, ** denotes *p* < 0.01, *** denotes *p* < 0.001).

| Trait | n | df | Sum Sq | Mean Sq | F-value | p | Significance |
| --- | --- | --- | --- | --- | --- | --- | --- |
| Juvenile Length ([log]mm at Day 7) | 83 |  |  |  |  |  |  |
| Treatment |  | 2 | 0.8830 | 0.4415 | 7.142 | 0.0014 | ** |
| Dose |  | 3 | 1.7520 | 0.5841 | 9.450 | 1.97E-05 | *** |
| Interaction |  | 3 | 0.6730 | 0.2244 | 3.631 | 0.0163 | * |
| Residuals |  |  | 5.0680 | 0.0618 |  |  |  |
| Adult Length ([log]mm at Day 21) | 62 |  |  |  |  |  |  |
| Treatment |  | 2 | 0.0222 | 0.0111 | 1.082 | 0.3431 |  |
| Dose |  | 3 | 0.1384 | 0.0461 | 4.493 | 0.0064 | ** |
| Interaction |  | 3 | 0.0910 | 0.0303 | 2.952 | 0.0394 | * |
| Residuals |  |  | 0.6368 | 0.0103 |  |  |  |
| Growth Rate (Adult Length - Juvenile Length<br>[log(mm)]/Juvenile Length [log(mm)]/Day) | 62 |  |  |  |  |  |  |
| Treatment |  | 2 | 0.1071 | 0.0536 | 1.323 | 0.2739 |  |
| Dose |  | 3 | 0.2817 | 0.0939 | 2.319 | 0.0841 |  |
| Interaction |  | 3 | 0.3887 | 0.1296 | 3.199 | 0.0294 | * |
| Residuals |  |  | 2.5108 | 0.0405 |  |  |  |
| Mass ([log]µg) | 46 |  |  |  |  |  |  |
| Treatment |  | 2 | 0.0521 | 0.0260 | 0.517 | 0.6000 |  |
| Dose |  | 3 | 0.1560 | 0.0520 | 1.032 | 0.3870 |  |
| Interaction |  | 3 | 0.2725 | 0.0908 | 1.803 | 0.1600 |  |
| Residuals |  |  | 2.1318 | 0.0504 |  |  |  |
| O2 Consumption (mg O2/hr/mg of dry mass) | 43 |  |  |  |  |  |  |
| Treatment |  | 2 | 0.0000 | 0.0000 | 0.122 | 0.8860 |  |
| Dose |  | 3 | 0.0000 | 0.0000 | 0.622 | 0.6050 |  |
| Interaction |  | 3 | 0.0000 | 0.0000 | 1.297 | 0.2880 |  |
| Residuals |  |  | 0.0002 | 0.0000 |  |  |  |

**Table 3.**
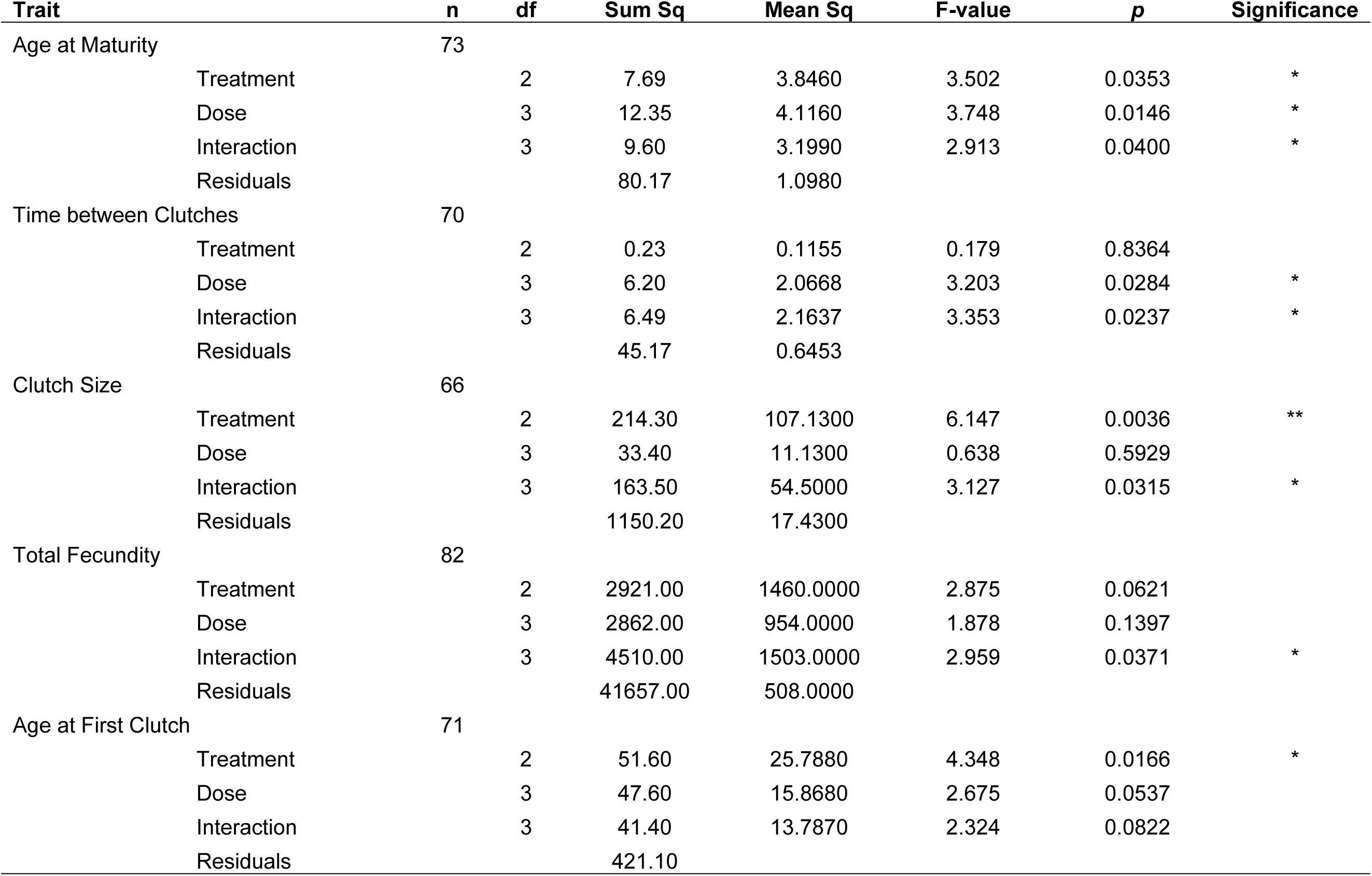
Effects of estrogenic compounds (estriol and EE2) across a range of doses (0, 0.1 µg/L, 1 µg/L, 10 µg/L, and 50 µg/L) on reproduction and fecundity traits for *D. magna* (* denotes *p* < 0.05, ** denotes *p* < 0.01, *** denotes *p* < 0.001).

### Estrogen concentration affects fitness traits

Overall, we observed effects of estrogen on fitness traits at certain concentrations of either EE2 and/or estriol when compared to unexposed control animals (Tables 1, 2, and 3). Survivorship curves for EE2 (Figure 2A), estriol (Figure 2B), and both estrogens combined (Figure 2C) illustrate survival differs depending on concentration. Notably, low concentrations of both EE2 and estriol increase the probability of survivorship, whereas high concentrations decrease the probability of survival, relative to the survival rates observed in control animals (depicted with a black line in all three panels of Figure 2).

For many other traits assayed, there was no simple positive or negative relationship between concentration and the traits assayed (Figure 3 and 4). Overall, dose was a significant main effect for five of the eleven fitness traits measured, and contributed to an interaction effect for seven traits (Tables 1, 2, and 3). In many cases, however, the animals experiencing the highest doses of the two estrogenic compounds did not exhibit the largest effects (e.g., effects on age at maturity; Figure 4A). In many cases, dosage effects differed depending on the type of estrogen, for example in the case of the effect of low doses on mean clutch size (Figure 4D).

**Figure 3.**
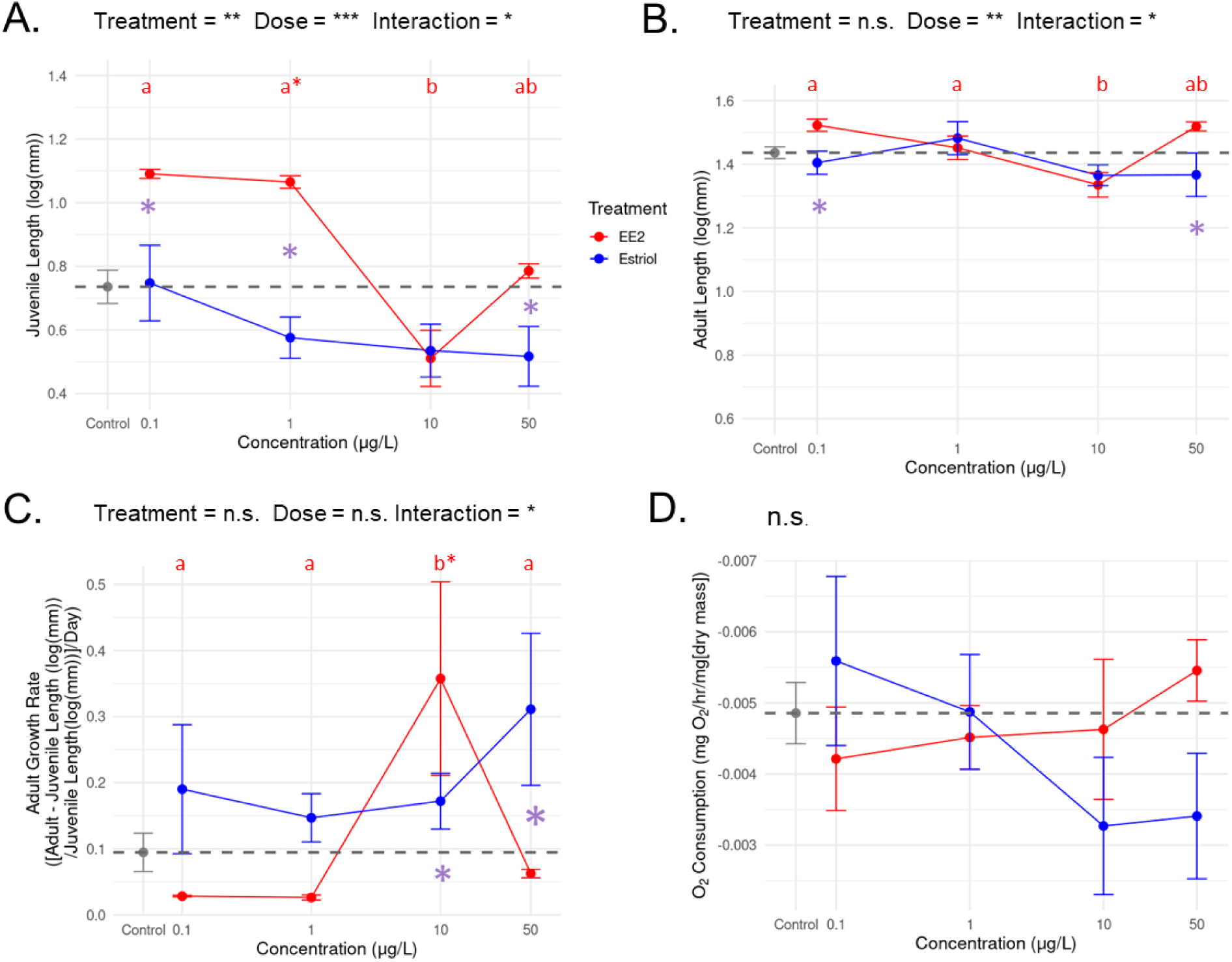
Mean values (±SE) of responses to estrogen exposures for four traits related to growth and metabolism, with main and interaction effects indicated above each pane. For each trait, 17α-ethinylestradiol (EE2) is plotted in red and estriol is in blue with the mean for the controls depicted by the gray dashed line. Differences between dosage levels are denoted with letters for each treatment and between estrogen types are denoted with an asterisk (purple). Cases where the mean for one type differed from the control, but not the other estrogen type, are depicted with a red (for EE2) or blue (for estriol) asterisk. Traits are (A) juvenile length (log[mm]) measured on day seven, (B) adult length measured on day twenty-one, (C) adult growth rate, and (D) oxygen (O_2_) consumption rates used to estimate metabolism.

### Estrogen type affects fitness traits and interaction effects are prevalent

Despite slight structural differences between estriol and EE2 (Figure 1), there are some major observable differences in the fitness consequences for animals exposed to these two forms. Overall, EE2 had more effects (impacting four traits, as opposed to three) and more variable effects (see lower case ‘a’ and ‘b’ designations of differences among dosage levels for each estrogen compound [red = EE2 and blue = estriol] in Figures 3 and 4). An interaction effect between the form of estrogen and dose was observed for seven out of the eleven traits assayed in our study (Tables 2 and 3), and in two cases (adult growth rate and total fecundity) this was the only significant effect, underscoring the importance of testing multiple concentrations when looking at the impact of hormones on fitness. In four traits, non-monotonic patterns were observed, where the relationship between dose and the trait values switch directions multiple times across the range of exposure levels (e.g., time between clutches; Figure 4C). While we observed an interaction effect on adult growth rates, our assay of oxygen consumption (a proxy for metabolic rate and respiration) revealed no significant differences. This could be due, however, to reduced sample sizes (n = 43) at the end of the exposure period or because oxygen consumption is not a sensitive enough indicator of differences in metabolic function.

**Figure 4.**
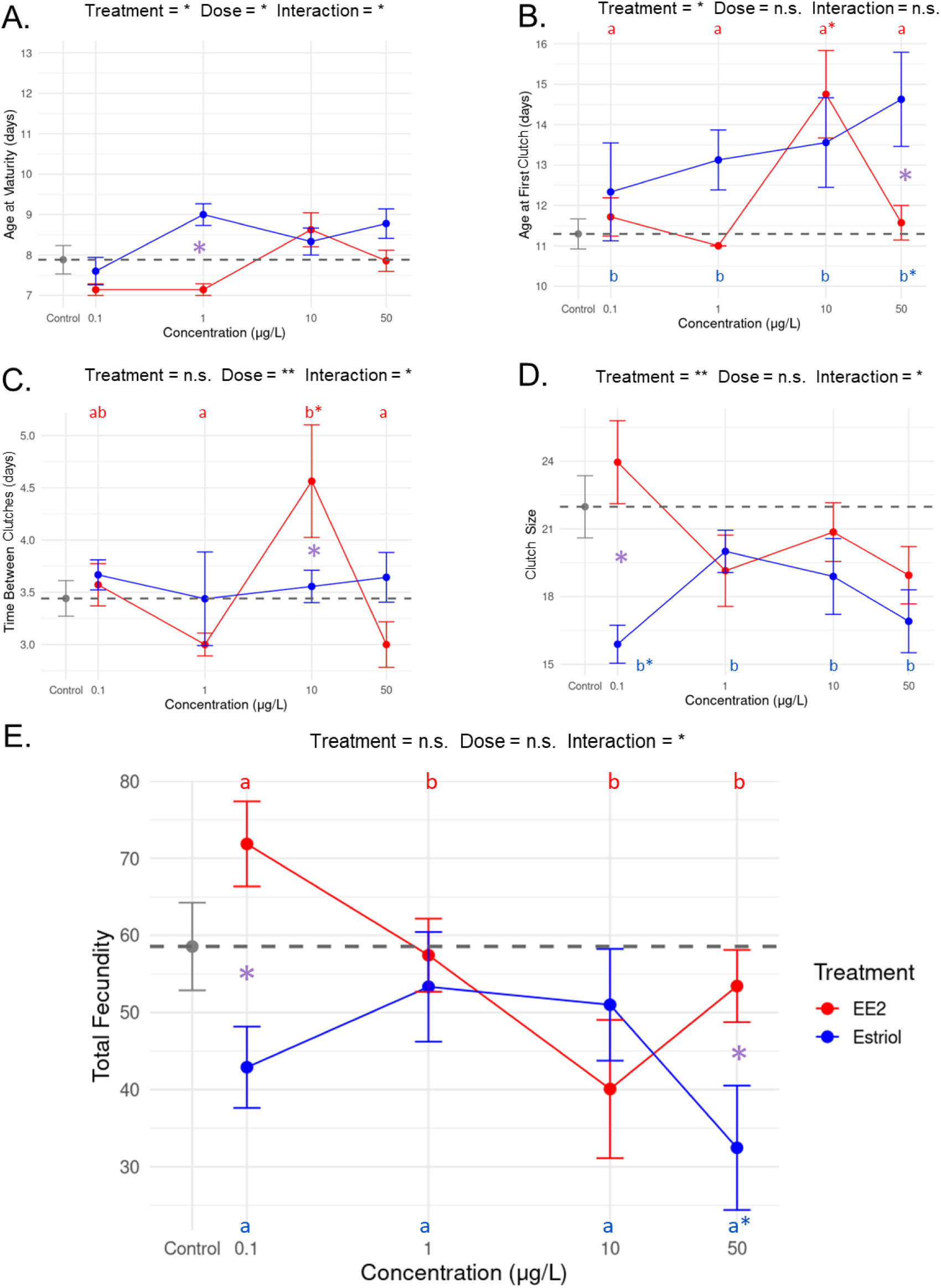
Mean values (±SE) of responses to estrogen exposures for traits related to reproduction and fecundity, with main and interaction effects indicated above each pane. For each trait, 17α-ethinylestradiol (EE2) is plotted in red and estriol is in blue and the mean for the controls is depicted by the gray dashed line. Significant differences between dosage levels are denoted with letters for each treatment and between estrogen types are denoted with an asterisk (purple). Cases where the mean for one type differed from the control, but not the other estrogen type, are depicted with a red (for EE2) or blue (for estriol) asterisk. Traits are (A) age at maturity (visible eggs in the brood chamber), (B) age at first clutch, (C) time between clutches, (D) clutch size, and (E) total fecundity.

## Conclusion

Taken together, our data support growing concern about the impact of estrogenic compounds in the environment, and specifically for the Daphnia as a key species in freshwater aquatic ecosystems. The effects of estrogen exposure vary based on both dose and type, in some cases opposing effects depending on dosage and type (as in Lange et al. 2012). Not all effects on fitness were negative, however. For example, changes in survival rates suggest that exposure to low levels of estrogen, at least for *Daphnia*, may be beneficial, whereas high concentrations strongly reduced survival (Figure 2). For traits related to growth and metabolism, it is clear that estrogen exposure can lead to increases or reductions in body size or growth rate for exposed animals, but the direction of the shifts depends on the form of estrogen. Finally, in the case of reproductive traits, such as age at maturity or total fecundity, again, both the form and dose of estrogen play a critical role in determining whether these traits lead to a net benefit or decline in fitness.

Our results have implications not only for *Daphnia*, but for the impact of estrogen pollution on entire aquatic ecosystems given the role they play in multitrophic cascades (Hallgren et al. 2014, Adeel et al. 2017). Future studies on the effects of estrogens should aim to expand the number and type of phenotypic assays (e.g., metabolomic function [O’Rourke et al. 2023]) and begin to investigate the mechanisms underlying such effects. Furthermore, long-term studies could shed light on the possibility of developing resistance with prolonged exposure (Clubbs and Brooks 2007). Estrogen pollution is likely to remain an issue, and removal methods are costly (including oxidation, photocatalysis, adsorption, and membrane filtration [Ghazal et al. 2022; Ben et al. 2017; Klaic and Jirsa 2022]). As a result, the current levels of estrogenic compounds encountered by organisms in their environment may be highly variable and/or increasing, thus comparing effects at multiple dosages remains a priority (Barrieros et al. 2016). Examining the variable phenotypic consequences of estrogen exposure in *Daphnia* can provide critical insights into the ways in which hormone pollution may shape our environment and well-being.

## Author Contributions

SJB and SS conceived of the study. SJB conducted the experiments and performed the assays. SJB and SS analyzed/interpreted the data and wrote the manuscript.

## Acknowledgments

We would like to thank Dieter Ebert for supplying the ancestral lineage of *D. magna* used in this study. We would also like to acknowledge funding from the M.J. Murdock Charitable Trust and the National Science Foundation (DEB-2336342) to SS.

## Supplemental Materials

Supplemental File S1. R code for all analyses.

Supplemental Table S1. Data table.

Supplemental Table S2. Model testing.

Supplemental Table S3. Post hoc tests for Fig 3.

Supplemental Table S4. Post hoc tests for Fig 4.

